# Multi-omic analysis of bat versus human fibroblasts reveals altered central metabolism

**DOI:** 10.1101/2023.05.08.537600

**Authors:** N. Suhas Jagannathan, Javier Yu Peng Koh, Younghwan Lee, Radoslaw M Sobota, Aaron Irving, Lin-Fa Wang, Yoko Itahana, Koji Itahana, Lisa Tucker-Kellogg

**Affiliations:** Cancer and Stem Cell Biology Programme, Duke-NUS Medical School, 8 College Road, Singapore, 169857, Singapore; Centre for Computational Biology, Duke-NUS Medical School, 8 College Road, Singapore, 169857, Singapore; Functional Proteomics Laboratory, Institute of Molecular and Cell Biology (IMCB), Agency for Science, Technology and Research, A*STAR, Singapore 138648, Singapore; Programme in Emerging Infectious Diseases, Duke-NUS Medical School, 8 College Road, Singapore, 169857, Singapore; Zhejiang University-University of Edinburgh Institute, Zhejiang University School of Medicine, Zhejiang University, Haining, China; SingHealth Duke-NUS Global Health Institute, Singapore, Singapore

**Keywords:** Bat metabolism, ferroptosis, ischemia, flux modeling, succinate accumulation

## Abstract

Bats have unique characteristics compared to other mammals, including increased longevity and higher resistance to cancer and infectious disease. While previous studies have analyzed the metabolic requirements for flight, it is still unclear how bat metabolism supports these unique features, and no study has integrated metabolomics, transcriptomics, and proteomics to characterize bat metabolism. In this work, we performed a multi-omics data analysis using a computational model of metabolic fluxes to identify fundamental differences in central metabolism between primary lung fibroblast cell lines from the black flying fox fruit bat (*Pteropus alecto*) and human. Bat cells showed higher expression levels of Complex I components of electron transport chain (ETC), but, remarkably, a lower rate of oxygen consumption (OCR). Computational modeling interpreted these results as indicating that Complex II activity may be low or reversed, similar to an ischemic state. An ischemic-like state of bats was also supported by decreased levels of central metabolites and increased ratios of succinate to fumarate in bat cells. Ischemic states tend to produce reactive oxygen species (ROS), which would be incompatible with the longevity of bats. However, bat cells had higher antioxidant reservoirs (higher total glutathione and higher ratio of NADPH to NADP) despite higher mitochondrial ROS levels. In addition, bat cells were more resistant to glucose deprivation and had increased resistance to ferroptosis, one of the characteristics of which is oxidative stress. Thus, our studies revealed distinct differences in the ETC regulation and metabolic stress responses between human and bat cells.

## Introduction

Bats display many characteristics that set them apart from other mammals, including the capacity for wing-powered flight, low rates of cancer incidence (Seluanov *et al*, 2018), high longevity quotient (Austad & Fischer, 1991; Austad, 2010; Wilkinson & South, 2002), and ability to carry many viruses (as a reservoir) without ill health (Wang *et al*, 2011). Each of these traits has distinct metabolic requirements and can affect overall metabolic activity. For example, flight is an energy-intensive process that requires high metabolic rates and ATP production (Thomas & Suthers, 1972; Maina, 2000). Resistance to cancer depends on multiple factors such as ROS management, DNA repair (Huang *et al*, 2016, 2019), and the ability to efflux genotoxic compounds (Koh *et al*, 2019), all of which have metabolic underpinnings. Longevity and disease resistance both require tolerance to metabolic and oxidative stresses, and the ability to dampen inflammasome activation (Ahn *et al*, 2019; Kacprzyk *et al*, 2017).

Multiple studies have documented individual aspects of metabolic regulation in bats. Metabolic rates of bats during flight (ATP production) have been documented to be approximately three times greater than basal metabolic rates in other mammals of similar size (Thomas & Suthers, 1972). In most other mammalian species, increased ATP production also creates increased production of reactive oxygen species (ROS) in the mitochondria, which can eventually lead to DNA and cellular damage (Buffenstein *et al*, 2008), and activate inflammasome responses. On the other hand, low amounts of ROS are a part of homeostasis and have been linked to beneficial effects on survival in multiple contexts (Mittler, 2017; Di Meo *et al*, 2016). It is conceivable that the homeostatic amount of ROS for a healthy cell is species-specific, as different species may have different ways of coping with the adverse effects of ROS accumulation, e.g., ROS-induced DNA damage and lipid peroxidation, or may activate downstream pathways such as inflammasomes at different levels of ROS. Hence for bats to be able to have a high metabolic rate without concomitant cellular/DNA damage would require one of the following to be true – improved decoupling of ATP production from ROS production (reduced leakage of electrons, so less ROS is produced), improved antioxidant defense to neutralize generated ROS, or improved repair of damage resulting in fewer deleterious effects of ROS (e.g., less inflammasome activation).

Different studies have proposed different mechanisms of ROS tolerance or antioxidant defense in bats including lower hydrogen peroxide production (Brunet-Rossinni, 2004; Brunet-Rossinni & Austad, 2004; Podlutsky *et al*, 2005; Ungvari *et al*, 2008), improved DNA repair (Foley *et al*, 2018), higher expression of heat shock proteins (Chionh *et al*, 2019) and/or a drug efflux factor, ABCB1 (Koh *et al*, 2019), and positive selection for efficient mitochondria. While the reasons could be multifactorial, no studies have performed a systems-wide comparison of mitochondrial metabolism between bats and higher mammals such as humans. Such characterization of the basal energy metabolism of bats and how it is different from other mammals such as humans, could shed light on mitochondrial activity/ROS management and hint toward metabolic factors that underlie/support desirable traits in bats, e.g., longevity, low cancer/mutation rates, and disease tolerance. Toward this goal, we compared the basal metabolism of cells from black flying fox fruit bat *Pteropus alecto* (*P. alecto*) and human cells.

*P. alecto* is a member of the pteropodidae family and is among the largest fructivore bats in the world. It has a lifespan of over 20 years and is documented to co-exist with lethal zoonotic viruses like Hendra and Nipah viruses. Using a primary lung fibroblast cell line that our group had established earlier from *P. alecto* (PaLung), we conducted a comparison between PaLung cells and human primary fibroblasts WI-38 cells to elucidate fundamental metabolic differences in the mitochondria.

Since bats and humans are very different species, it is likely that data from any one high-throughput platform (e.g., transcriptomics) would show many differences. Hence, to identify consistent differences in the metabolic regulation of *P. alecto* and humans, we looked for concordance between multiple high-throughput platforms: whole-cell transcriptomics, mitochondrial proteomics, and whole-cell metabolomics. To integrate the different omics results, we use constraint-based flux sampling, which is a computational modeling technique that simulates metabolic flux patterns using existing knowledge about metabolic network connectivity and topology. Constraint-based flux modeling has been used previously for comparing metabolic phenotypes across cancers (Aurich *et al*, 2017), understanding metabolic regulation of macrophage polarization (Bordbar *et al*, 2012), studying ischemia-reperfusion injury (Chouchani *et al*, 2014), characterizing microbiomes (Jansma & El Aidy, 2021; Ezzamouri *et al*, 2023), optimizing metabolite production (Patil *et al*, 2004), and understanding metabolic contributors of disease pathology e.g., diabetes (Ravi & Gunawan, 2021).

Here, our results show that PaLung cells have differences in basal metabolism that resemble ischemia, associated with the low or reverse activity of Complex II in the electron transport chain (ETC). Finally, we confirmed our prediction of ischemic-like basal metabolism in PaLung cells by characterizing the response of bat cells to cellular stresses such as oxidative stress, nutrient deprivation, and a type of cell death related to ischemia, *viz.* ferroptosis.

## Results

### Transcriptomics identifies differences in oxidative phosphorylation between PaLung cells and WI-38 cells

To understand the differences in cellular-scale metabolism between PaLung cells (*P. alecto*) and WI-38 cells (*H. sapiens*), we performed whole-cell transcriptomics on the two cell lines. Transcriptomics detected a total of 21,952 mRNA transcripts in bat PaLung cells and 58,830 transcripts in human WI-38 cells, respectively. Since the bat genome is not as well annotated as the human genome, we performed downstream differential expression analysis using the set of 14,986 common transcripts found in both PaLung and WI-38 cells. Differentially expressed genes were detected using the DEseq pipeline, which was modified to normalize for the different transcript lengths in both species (see Methods, Figure 1A, Supplementary Figure S1A). This method yielded a total of 6,247 differentially expressed (DE) transcripts (|log fold change| >= 1 and FDR < 0.05, Figure 1B, Supplementary Figure S1B). Because there is no standard way of normalizing RNAseq data for inter-species comparison, we also repeated the analysis in the EdgeR package, using a recently published normalization method GeTMM (Smid *et al*, 2018). Both methods yielded very similar results with minor discrepancies (Supplementary Figure S1C), and we chose to perform further downstream analysis using the DEseq-generated DE transcript list. A summary of transcriptomics results for core metabolic pathways can be found in Supplementary Figure S2, and the list of differentially expressed genes can be found in Supplementary Table 1. The number of differentially expressed genes (6,247) was extremely high, suggesting that multiple pathways are differentially regulated between the two species. Since our primary goal is to understand species-specific differences in metabolism, we filtered our gene dataset to contain only genes listed under the Gene ontology category Cellular Metabolic Process (GO ID:0044237), resulting in a truncated list of 4794 genes. We then performed gene set enrichment analysis (GSEA) using the expression of these 4794 metabolic genes as input, searching against the Gene Ontology Biological Process (GO-BP) database. Supplementary Tables 2 and 3 contain the list of enriched gene sets in PaLung and WI-38 cells respectively. Two of the top differentially-regulated metabolic gene sets identified by GSEA were (a) Respiratory electron transport, ATP synthesis by chemiosmotic coupling, and heat production by uncoupling proteins, and (b) Cellular response to hypoxia, in the Reactome Pathway Database. The genes belonging to both oxidative phosphorylation (OxPhos) (Figure 1C) and response to hypoxia (Figure 1D) gene sets had increased transcriptional expression in PaLung cells. This was interesting because, conventionally, these two pathways are active under opposing conditions (high oxygen for OxPhos vs low oxygen for hypoxia), and so would not be expected to vary in concert with each other. This hinted toward non-trivial regulation of central metabolism in PaLung cells.

**Figure 1.**
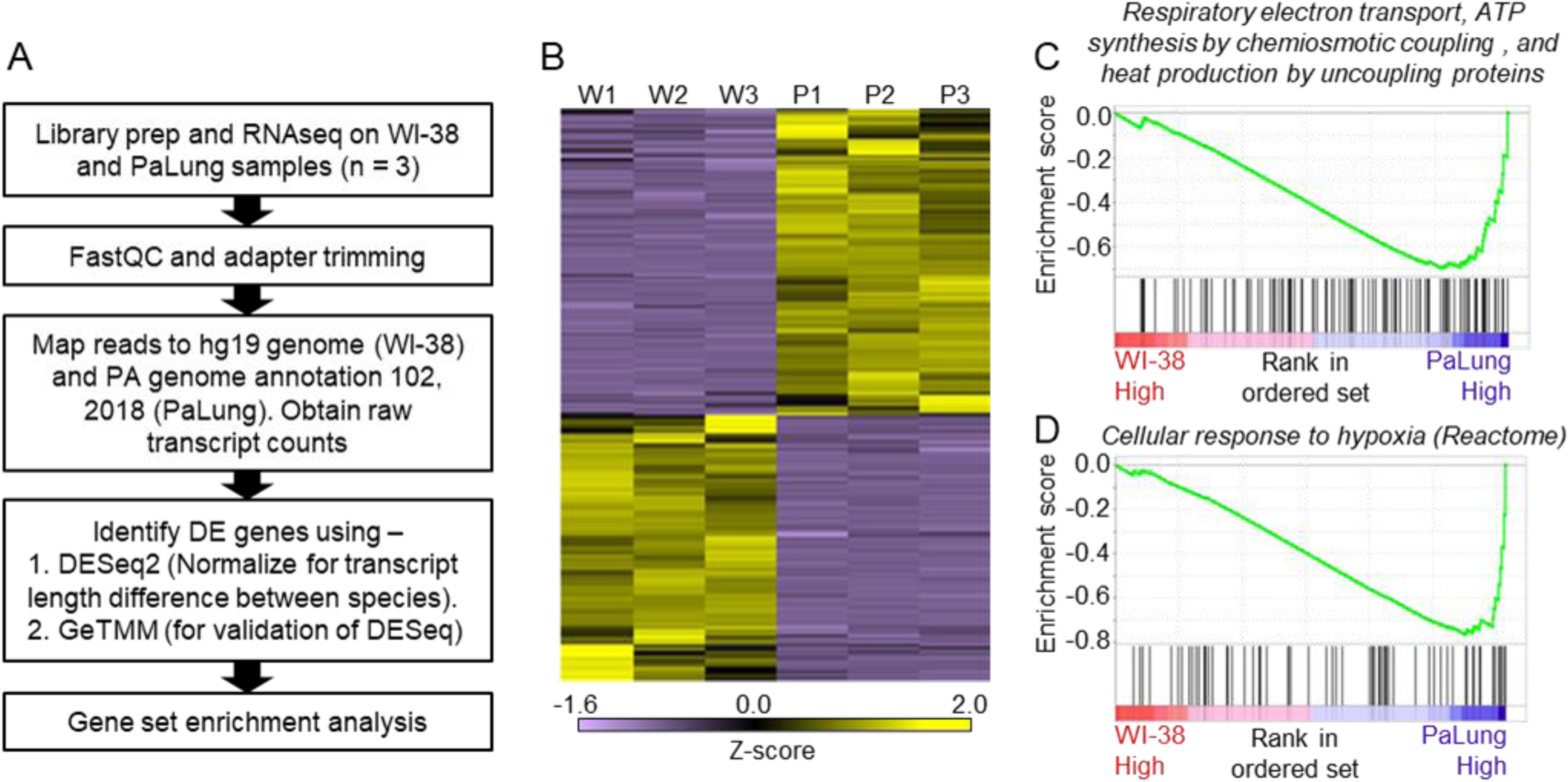
RNAseq data analysis of PaLung and WI-38 cells for differential expression and pathway enrichment. (A) Workflow of bioinformatics analysis pipeline for RNAseq data from PaLung (P. alecto) and WI-38 (H. sapiens) cells (n = 3). (B) Heatmap showing the expression patterns for differentially expressed genes in the three WI-38 samples (W1-W3) and the three PaLung samples (P1-P3). (C-D) GSEA analysis identifies Respiratory electron transport and Cellular response to hypoxia as top metabolic pathways that are differentially regulated between PaLung and WI-38 cells. Shown here are the enrichment score plots for (C) respiratory electron transport and (D) cellular response to hypoxia.

### Mitochondrial Proteomics suggests that oxidative phosphorylation is higher in PaLung cells compared to WI-38 cells

To test whether the whole-cell RNA differences were also reflected in mitochondrial composition, we performed tandem mass tag-based proteomic profiling in the mitochondrial fractions of PaLung and WI-38 cells. Profiling detected a total of 1,469 proteins. Analysis using the gene ontology tool Enrichr confirmed that a majority of these proteins were likely obtained from a mitochondrial compartment (Supplementary Figure S3A). When we performed differential expression analysis on the 1469 proteins (see Methods, Figure 2A), 405 were differentially expressed between WI-38 and PaLung cells (Supplementary Figure S3B). Of these 405 proteins, we identified 127 to be core mitochondrial proteins (as defined by Mitocarta and IMPI datasets), that were differentially expressed between WI-38 and PaLung cells (Supplementary Table 4). Figure 2B shows the heatmap of row-normalized abundances of the 127 DE proteins. We observed that most of these 127 DE mitochondrial proteins are upregulated in bat PaLung samples (109), with very few downregulated proteins (18), suggesting increased mitochondrial activity in PaLung cells.

**Figure 2.**
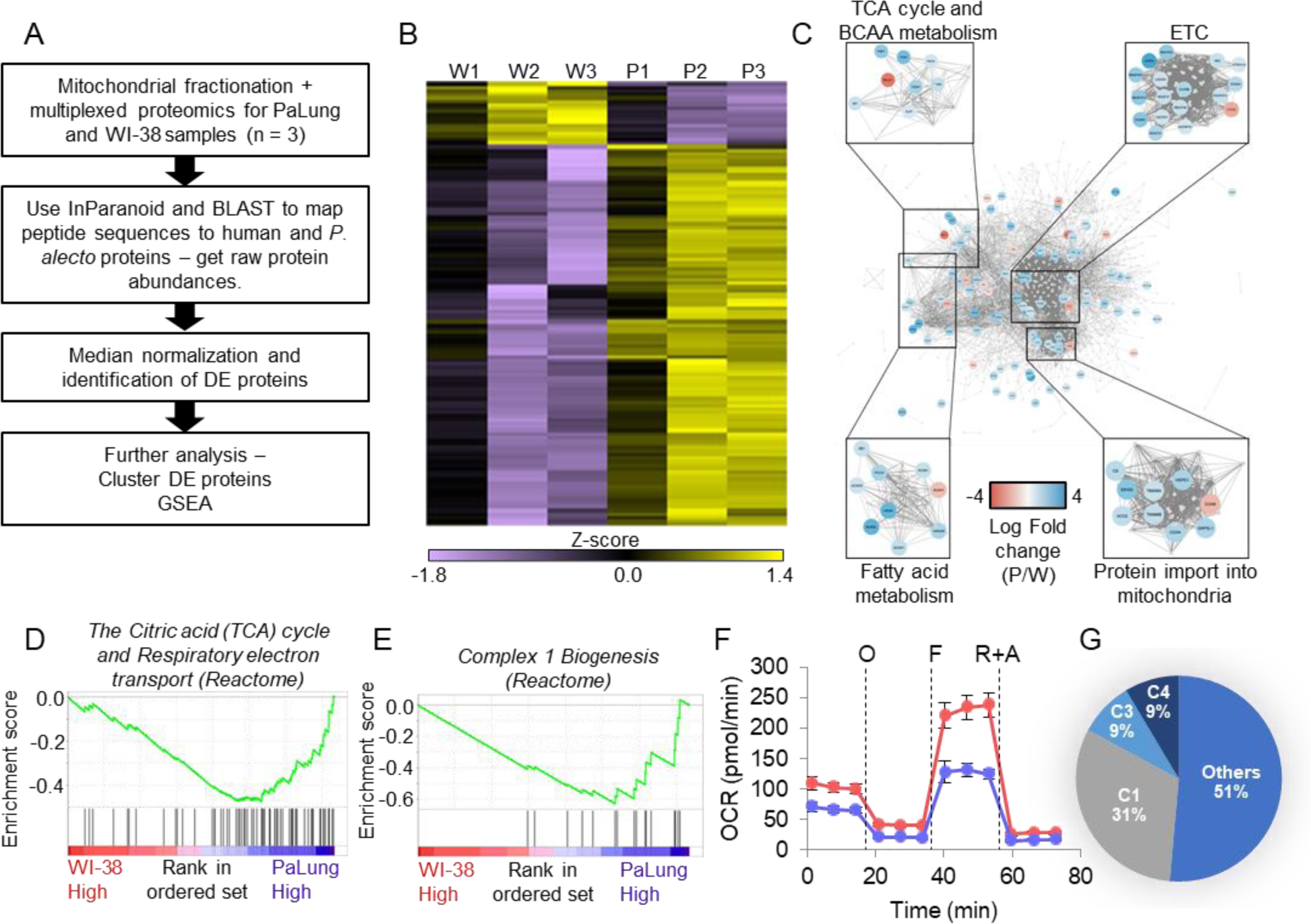
Proteomic data analysis of mitochondrial fractions of PaLung and WI-38 cells for differential expression, pathway enrichment, and ETC activity. (A) Workflow of bioinformatics analysis pipeline for proteomics data from PaLung (*P. alecto*) and WI-38 (*H. sapiens*) cells (n = 3). (B) Heatmap showing the expression patterns for the 129 differentially expressed mitochondrial proteins in the three WI-38 samples (W1-W3) and the three PaLung samples (P1-P3). (C) Differentially expressed mitochondrial proteins (nodes colored by log fold change) are overlaid on a network of mitochondrial protein-protein interactions (obtained from STRING) (W = WI-38 cells, P = PaLung cells). The nodes are then clustered with respect to Reactome-annotated pathways. (D-E) GSEA analysis identifies Citric acid cycle, oxidative phosphorylation and Complex I biogenesis as top metabolic pathways that are differentially regulated between PaLung and WI-38 cells. Shown here are the enrichment score plots for (D) citric acid cycle and oxidative phosphorylation and (E) Complex I biogenesis. (F) OCR measurement of PaLung cells (blue) and WI-38 cells (red) plotted as mean ± SD from n > 15 independent experiments. O = oligomycin, F = FCCP, R+A = rotenone+antimycin A. (G) Pie chart showing that the proteomic upregulation of the ETC, implied by GSEA analysis, is dominated by genes for subunits of Complex I. C1 = Complex I; C3 = Complex III; C4 = Complex IV of the ETC.

To identify highly connected subnetworks that have strong expression changes, the set of 127 DE proteins was overlaid on a background of all known mitochondrial protein-protein interactions using STRING (with background obtained from Mitocarta and IMPI datasets). The results were clustered, and Reactome pathway enrichment analysis of the resulting clusters showed enrichment for the following pathways: (1) TCA cycle and BCAA metabolism, (2) electron transport chain (3) fatty acid metabolism, and (4) protein import into mitochondria (Figure 2C).

We also performed GSEA using the abundances of the 1469 detected proteins as input against gene sets in the GO BP database. Supplementary Tables 5-6 show the enriched gene sets in PaLung cells and WI-38 cells respectively. Supplementary Table 5 shows that PaLung mitochondria were enriched for proteins in respiratory electron transport compared to WI-38 cells, (Figure 2D). In particular, the gene set for Complex I biogenesis (Figure 2E) was significantly enriched in PaLung cells. Proteomics thus agrees with transcriptomics that OxPhos genes are more highly expressed in PaLung cells than WI-38 cells, however, there were no indications of hypoxia-related gene sets being differentially regulated in the proteomics dataset. Hence, we decided to pursue experimental studies of OxPhos rates in PaLung and WI-38 cells.

### PaLung cells have a lower oxygen consumption rate (OCR) than WI-38 cells

To better characterize differences in OxPhos between PaLung and WI-38 cells, we monitored the oxygen consumption of these two cell lines using Seahorse XF Analyzer. Unexpectedly, the oxygen consumption rate (OCR) under basal conditions was lower in PaLung cells compared to WI-38 cells. Notably, bat cells have much lower maximal respiratory capacity compared to WI-38 cells, indicated by FCCP treatment (Figure 2F). This suggested that PaLung cells are able to carry out less oxidative phosphorylation than WI-38 cells. While this observation agrees with earlier studies showing mild-depolarization in the mitochondria of bats (Vyssokikh *et al*, 2020), it is in sharp contrast with our observation of increased OxPhos machinery in PaLung cells, using transcriptomics and proteomics. Taking a closer look at the omics results, we observed that the GSEA-flagged upregulation in OxPhos was driven mostly by the upregulation of Complex I subunits, for both the proteomic and transcriptomic data (Figure 2G, Supplementary Figure S1D). This led us to hypothesize that in the basal state, PaLung cells might have partial decoupling of Complex I from the electron transport chain (ETC), meaning that electrons emerging from Complex I might not proceed through the entirety of the ETC. This partial decoupling of Complex I might result in lower overall ATP synthesis, creating continued demand for ATP production from other sources. This would be consistent with our transcriptomic finding that both OxPhos-related and hypoxia-related genes were upregulated in PaLung cells. To build a self-consistent interpretation of these paradoxical omics and functional datasets, we proceeded to perform computational modeling of mitochondrial metabolism.

### Computational flux modeling suggests that Complex II of the ETC may run in reverse in PaLung cells

To understand the metabolic consequences of having higher Complex I activity but lower overall respiration, we turned to constraint-based flux modeling (Supplementary Figure S4). We started with Mitocore, a published model of mitochondrial metabolism, that includes both core mitochondrial reactions and supporting cytoplasmic reactions from central carbon metabolism (glycolysis, pentose phosphate pathway, folate cycle, urea cycle, etc.) (Smith *et al*, 2017). Mitocore provides the set of metabolic reactions that can occur in a cell without specifying the activity, abundance, expression, or utilization of each element. To establish a model for each species, we took a species-specific subset of Mitocore reactions (called a *context-specific reconstruction*) based on the presence and absence of gene/protein expression in the transcriptomic/proteomic data for each species (See Methods). This resulted in a PaLung model with 409 reactions and 324 metabolites (Supplementary Table 7), and a WI-38 model with 437 reactions and 341 metabolites (Supplementary Table 8) as shown in Figure 3A. We then performed uniform flux sampling (5,000 samples) of the two models without imposing any constraints on either model (Figure 3B). Simulations resulted in a feasible flux distribution for each reaction in both the PaLung model and the WI-38 model, which can be depicted as a frequency histogram. In each chart in Figure 3B and 3C, the X-axis represents the metabolic flux value for the corresponding reaction, and Y-axis represents the frequency/probability of the reaction having the specific flux value. Figure 3B shows these histograms for the fluxes through Complex I-III in both the PaLung and WI-38 metabolic models, under unconstrained conditions (control simulation). We next imposed the following two constraints, derived from the above transcriptomics/proteomics and oxygen consumption measurement. The first constraint was for the PaLung model to have higher activity of Complex I than the WI-38 model. The idea of this constraint is based on our transcriptomic/proteomic data (Figures 1C and 2F) into an assumption that the flux through Complex I in PaLung cells must be greater than the flux through Complex I in WI-38 cells (See “Constraints for flux sampling” in Methods). Based on our oxygen consumption data, the second constraint was for the WI-38 model to have higher oxygen intake into the mitochondria than the PaLung model. When the PaLung and WI-38 models were simulated under these constraints, the results revealed that the flux histograms for PaLung cells had shifted to very low or negative flux values for Complex II (Supplementary Table 9). Complex II is also called succinate dehydrogenase (SDH) and is part of both the TCA cycle and the ETC). SDH generally catalyzes the conversion of succinate into fumarate, accompanied by a reduction of the endogenous Quinone pool. However, SDH has also been documented to catalyze the reverse reaction, converting fumarate to succinate, although this is unconventional. Such an unconventional ETC paradigm where Complex I proceeds forward but Complex II proceeds in reverse (toward the accumulation of succinate) has been observed in other systems, e.g., in murine retinal tissues (Bisbach *et al*, 2020) and human hearts (Chouchani *et al*, 2014), under ischemic or hypoxic conditions. In these cases, the resulting accumulation of succinate was found to be useful in fueling other cells in the tissue or in avoiding reperfusion injury when ischemic conditions were abruptly removed. The low or negative flux values for Complex II in our PaLung simulations indicate that the electrons obtained from Complex I may accumulate at Complex II or potentially even get consumed by Complex II operating in reverse (bypassing the rest of the ETC) in PaLung cells. To further interrogate central carbon metabolism in PaLung cells and to validate if our predictions of unconventional SDH activity might be borne out by metabolic measurements, we undertook targeted metabolomics of small organic compounds for both PaLung and WI-38 cells.

**Figure 3.**
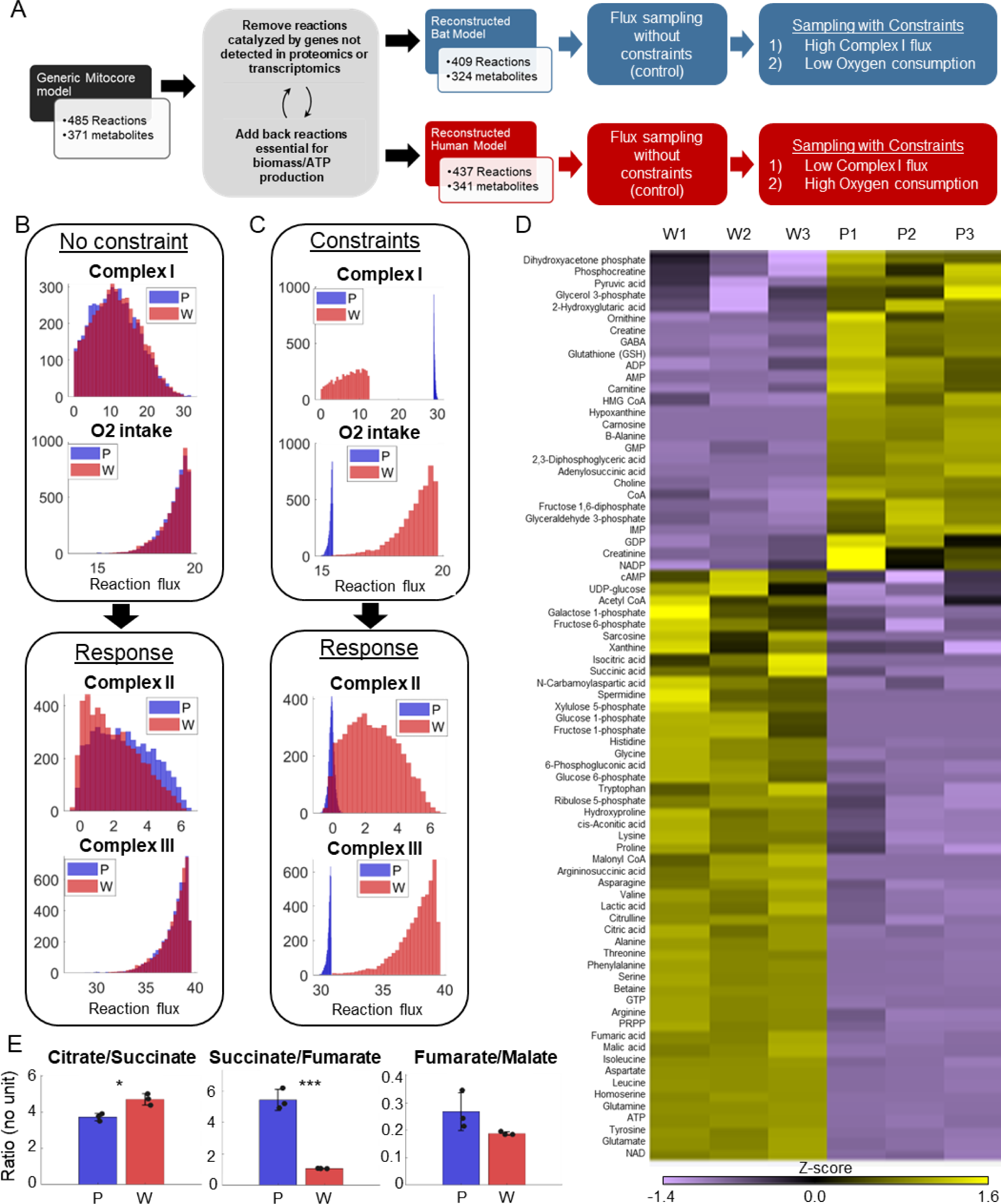
Metabolomic data and model-based analysis of mitochondrial metabolism in PaLung and WI-38 cells. (A) Schematic showing the metabolic modeling pipeline. We begin with the context-specific reconstruction of a metabolic model – the process where a generic mitochondrial model (Smith *et al*, 2017) is tailored specifically to PaLung and WI-38 cells using proteomic and transcriptomic expression patterns. The individual metabolic models are then simulated using constraint-based flux sampling methods to give a distribution of possible fluxes for each reaction in the model. Comparing these flux distributions between the two models allows the detection of metabolic reactions that are likely to be differentially regulated in response to user-imposed constraints on metabolism. Simulations are performed on both the PaLung and WI-38 models under no constraints or with constraints on Complex I and mitochondrial O2 intake. (B) Sample histograms showing the feasible flux distributions for ETC reactions in the unconstrained PaLung (P) and WI-38 (W) metabolic models. (C) Flux distributions of ETC reactions in PaLung (P) and WI-38 (W) cells when PaLung cells are constrained to have higher flux through Complex I of ETC but lower oxygen intake in the mitochondria. (D) Heatmap showing differentially regulated metabolites from central carbon metabolism in the three WI-38 (W1-W3) and the three PaLung (P1-P3) samples. (E) Intra-sample ratios of metabolites from the TCA cycle in either PaLung (P) or WI-38 (W) cells, plotted as mean ± SD from three independent experiments. * and *** represent p-value ≤0.05 or ≤0.001 respectively (unpaired Student’s two-sided t test with Benjamini-Hochberg correction for multiple hypothesis testing).

### Metabolomics supports low/reverse Complex II activity in PaLung cells

Mass spectrometry-based targeted metabolomics was performed on both PaLung and WI-38 cell lines, resulting in the absolute quantification of 116 metabolites from central carbon metabolism (Figure 3D, Supplementary Figure S5, Supplementary Table 10). Since we were comparing different species, we looked at relative ratios of metabolites within each species. As predicted by our hypothesis, metabolomics showed that the ratio of succinate-to-fumarate was much higher in PaLung cells (5.44 ± 0.67) compared to WI-38 cells (1.07 ± 0.012) (*p* < 0.001), consistent with succinate accumulation in PaLung cells (Figure 3E). In contrast, the ratio of other serial TCA metabolites e.g, citrate/succinate or fumarate/malate showed only mild or insignificant differences between PaLung and WI-38 cells. We interpreted this as an indication that the TCA cycle acts in a truncated manner in PaLung cells, with SDH operating in reverse to support succinate accumulation. Interestingly, previous studies have documented such SDH phenomena (with the low or reverse activity of Complex II) during ischemic/hypoxic states that had low metabolic rates and high levels of AMP (Chouchani *et al*, 2014; Bisbach *et al*, 2020). This led us to wonder if PaLung cell metabolism might resemble an ischemic-like state despite oxygen and nutrient availability, in which case we would expect to see low metabolic rates and high AMP levels in PaLung cells.

### PaLung cells exhibit basal metabolism that resembles an ischemic-like state and can tolerate glucose deprivation better than WI-38 cells

To further understand the consequences of the truncated TCA cycle and its implications on an ischemic-like state, we looked at the abundances of other metabolites in metabolomics datasets. Looking at metabolites from glycolysis, pentose phosphate pathway, TCA cycle, and the levels of amino acids, we found that most metabolites were at much lower levels in PaLung cells compared to WI-38 cells, except for a small subset of metabolites in glycolysis (Figure 4A-C). This suggested an overall slower metabolic turnover in PaLung cells. PaLung cells also had a much higher proportion of AMP (Figure 4D), which would be expected in an ischemic setting.

**Figure 4.**
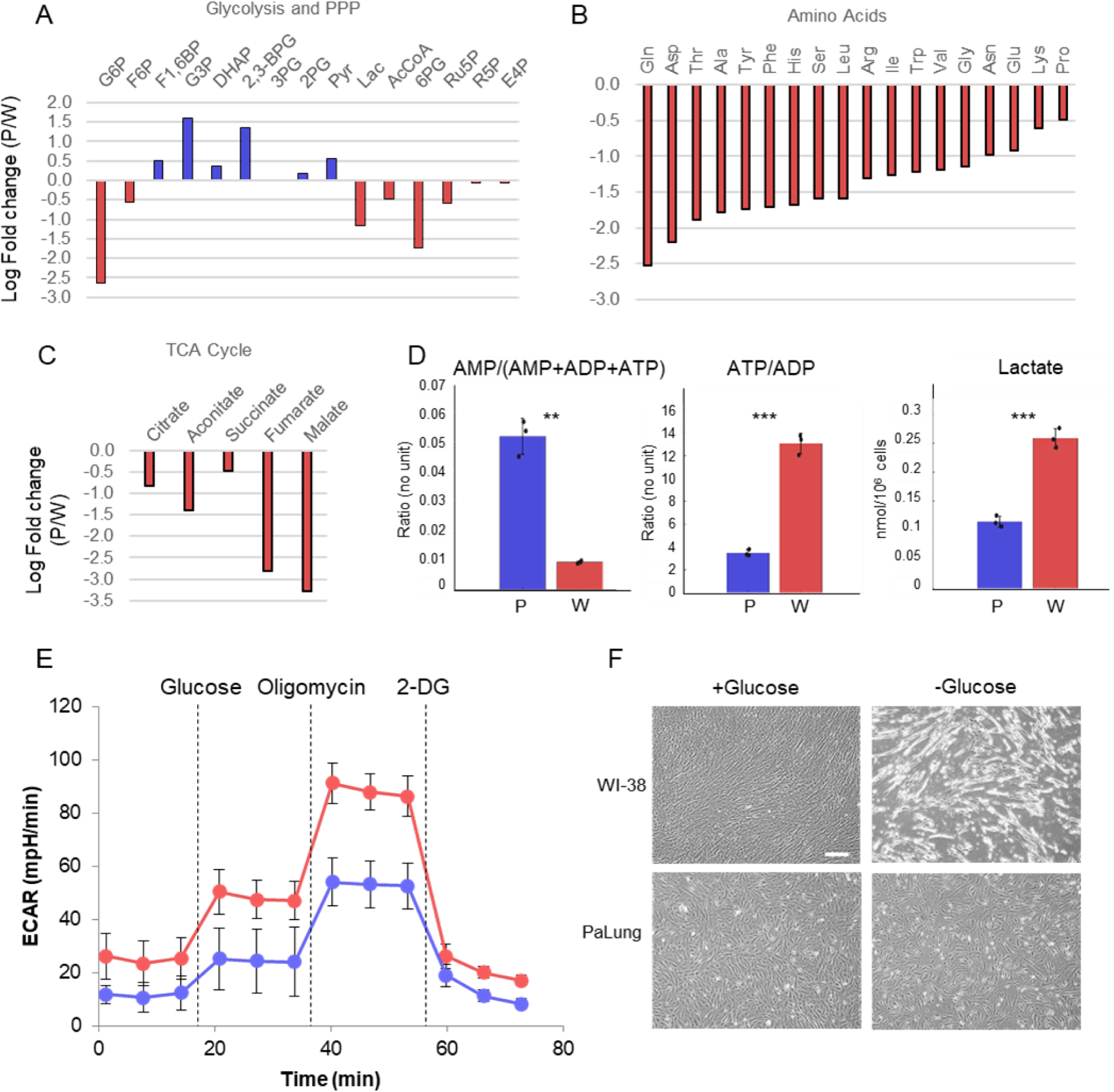
PaLung cells show basal metabolism that resembles an ischemic-like state. (A) Fold changes (mean PaLung/mean WI-38) of metabolite abundances, obtained through targeted metabolomics profiling of central carbon metabolites in PaLung and WI-38 cells. (B-C) Amino acid and metabolite changes in TCA cycle (PaLung/WI-38) in PaLung cells and WI-38 cells. (D) Bar plots of AMP/(AMP+ADP+ATP), ATP/ADP ratio, and lactate amounts in PaLung (P) and WI-38 (W) cells plotted as mean ± SD from three independent experiments. ** and *** represent p-value ≤0.01 or ≤0.001 respectively (unpaired Student’s two-sided t test with Benjamini-Hochberg correction for multiple hypothesis testing). (E) ECAR measurement of PaLung (blue) and WI-38 (red) cells plotted as mean ± SD from n > 15 independent experiments. (2-DG = 2-deoxy glucose). (F) Phase contrast images of PaLung and WI-38 cells with or without glucose deprivation for 96hrs. Scale bar 100 µm.

Furthermore, our metabolomics dataset revealed that PaLung cells had a 3-fold lower ATP/ADP ratio compared to WI-38 cells (*p* < 0.001). ATP/ADP ratio is a golden standard for the measurement of cellular energy status, and our results suggest lower metabolism in PaLung cells. Another corroborating measurement for energy status is adenylate energy charge, with lower values indicating slower metabolism (Atkinson & Walton, 1967). We found that PaLung cells had lower adenylate energy charge compared to WI-38 cells (*p* < 0.001) (Supplementary Table 10). From these parameters, we infer that PaLung cells have lower ATP synthesis, consistent with our earlier observations of lower OCR in PaLung cells (Figure 2F). To check if slower metabolism would also translate to slower glycolysis in PaLung cells, we first checked our transcriptomics and metabolomics data. Both omics datasets showed mixed signals along the glycolysis pathway (with partial upregulation and partial downregulation) (Figure 4A and Supplementary Figure S2), yielding no predictions for glycolysis utilization. However, metabolomics showed higher levels of intracellular lactate in WI-38 cells than PaLung cells, suggesting that PaLung cells could have lower glycolytic flux than WI-38 cells. Finally, we assessed glycolysis levels using a Seahorse XF Analyzer to quantify the extracellular acidification rate (ECAR). This revealed that PaLung cells had lower ECAR compared to WI-38 cells (Figure 4E). From these observations of low ECAR, low lactate, and low ATP/ADP ratio, combined with the earlier findings of downregulated OxPhos expression and low OCR, we conclude that PaLung cells have less energy production and a lower level of basal metabolic activity than WI-38 cells.

Reduced OxPhos and glycolysis in PaLung cells somewhat resemble an ischemic-like state with inadequate oxygen and glucose supply. Therefore, PaLung cells may be resistant to metabolic stress. To test this, we subjected PaLung and WI-38 cells to glucose deprivation for 96 hours. Interestingly, PaLung cells displayed higher viability than WI-38 cells after glucose deprivation (Figure 4F), despite starting with lower stores of internal energy (e.g., glucose-6-phosphate, acetyl-coA, and lower adenylate charge). These data suggest that PaLung cells do have a better ability to tolerate metabolic stress compared with WI-38 cells, consistent with low basal metabolic activity.

### PaLung cells have higher ROS compared to WI-38 cells but lower expression of many antioxidant genes

Previous studies of ischemic metabolism, in which SDH functions in reverse, have showed that high levels of ROS were generated as a result. To test if the same would be true in PaLung cells, we performed the mitosox assay to measure superoxide levels in the mitochondria of PaLung cells and WI-38 cells. Indeed, PaLung cells showed higher mitochondrial superoxide levels compared to WI-38 cells, both under basal conditions and when ETC was inhibited using antimycin A (Figure 5A). Searching our transcriptomic data for genes involved in the redox control, we observed that there were significant differences in the expression of redox control genes in PaLung and WI-38 cells. (Figure 5B). Notably, SOD1 and SOD2, key enzymes that convert mitochondrial superoxide to the more toxic intracellular ROS, are less expressed in PaLung cells than in WI-38 cells, consistent with higher mitochondrial superoxide levels in PaLung cells. Interestingly, glutathione peroxidase 3 (*GPX3*), a well-known antioxidant enzyme that reduces hydrogen peroxide or organic hydroperoxides using glutathione, was found to be highly upregulated in PaLung cells compared to WI-38 cells. This led us to test levels of glutathione, a non-enzyme antioxidant for ROS detoxification, in the two cell lines.

**Figure 5.**
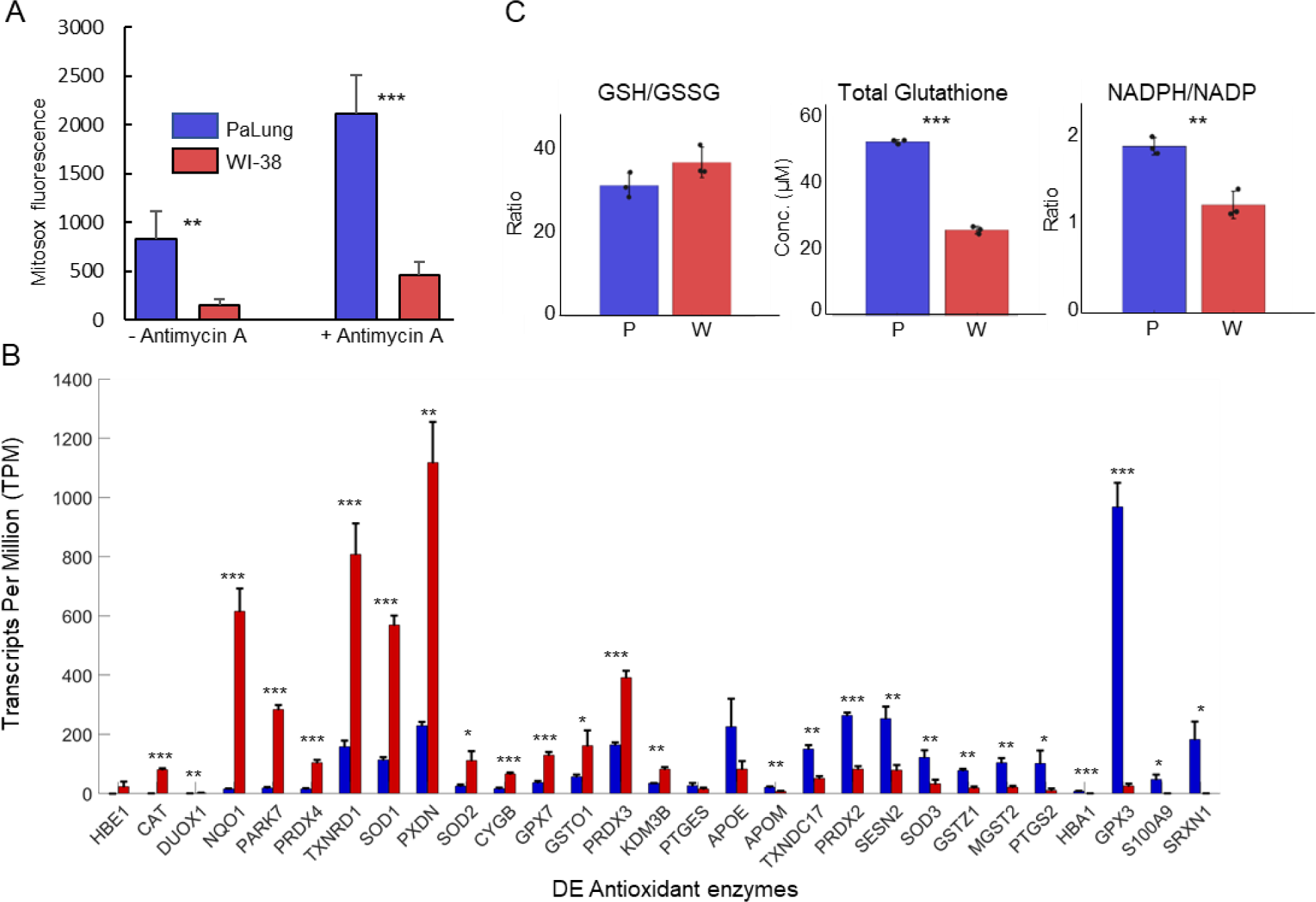
PaLung cells have higher ROS levels and lower antioxidant response than WI-38 cells. (A) MitoSOX measurement of PaLung and WI-38 cells with or without antimycin A treatment for 1 hr. Antimycin A is an ETC inhibitor known to induce superoxide generation. (B) Bar charts showing the expression levels of differentially expressed antioxidant genes (as transcripts per million, TPM) in PaLung (blue) and WI-38 (red) cells. Genes have been sorted in increasing order of P/W fold change. (C) Bar plots show the ratio of reduced to oxidized glutathione (GSH/GSSG), total Glutathione (GSH + GSSG), and the ratio of NADPH/NADP in PaLung (P) and WI-38 (W) cells. For all panels, bars are the mean ± SD from three independent experiments (n = 3). *, ** or *** represents p-value <0.05, ≤0.01, or ≤0.001 respectively (unpaired Student’s two-sided t test with Benjamini-Hochberg correction for multiple hypothesis testing).

### PaLung cells have a robust glutathione NADPH system to counter intracellular ROS

To test if the glutathione system may help PaLung cells tolerate intracellular ROS, we measured intracellular glutathione and NADP(H) levels in both PaLung and WI-38 cells. The ratio of reduced to oxidized glutathione (GSH/GSSG ratio) was not significantly different between the two cell lines (WI-38 cells: 36.45±3.6 vs PaLung cells: 30.94±2.97). However, PaLung cells had a two-fold higher concentration of total intracellular glutathione, compared to WI-38 cells (Figure 5C). We also found that PaLung cells had a higher NADPH/NADP ratio (1.5-fold) compared to WI-38 cells. Overall, the metabolomics data indicated that PaLung cells had a nearly 2.5-fold higher ratio of NADP/NAD, i.e., a higher resting concentration of phosphorylated to unphosphorylated NAD (Supplementary Figure S6A). This was also supported by transcriptomics which showed an upregulation of NADK (NAD kinase) and supporting enzymes required to synthesize and phosphorylate NAD (Supplementary Figure S6B). Taken together these results suggest that PaLung cells maintain a higher standing pool of glutathione and a higher NADPH concentration to counter ROS generated due to ischemic-like metabolism.

### PaLung cells are resistant to ferroptosis

Many earlier studies have shown that ischemic conditions can induce cell death via ferroptosis, which also depends on accumulated ROS and low glutathione levels (Chen *et al*, 2021). We wondered if apart from combatting ROS, the high glutathione levels might also help PaLung cells avoid ischemia-induced ferroptosis. Recent studies have reported that ischemia-induced ferroptosis causes tissue damage and that inhibition of ferroptosis attenuates ischemia-induced cell death (Xie *et al*, 2019; Liao *et al*, 2021). Ferroptosis is a non-apoptotic, programmed form of cell death, which is iron-dependent and occurs via glutathione depletion-induced lipid peroxidation (Ursini & Maiorino, 2020). Given that PaLung cells showed upregulation of glutathione/NADPH antioxidant system and genes related to hypoxia response (Figure 1D) and ischemic metabolism, we tested whether PaLung cells can better tolerate ferroptosis-inducing conditions. Ferroptosis was induced by erastin, an SLC7A11/xCT inhibitor. While WI-38 cells showed high sensitivity to erastin-induced ferroptosis that were prevented by a ferroptosis inhibitor ferrostatin-1, PaLung cells were resistant to erastin-induced ferroptosis (Figure 6A and B). Similarly, PaLung cells were more resistant to cystine deprivation-induced ferroptosis, compared to WI-38 cells and had almost 7-fold lower cell death than WI-38 cells (3.5% cell death in PaLung cells compared to 24.4% cell death in WI-38 cells upon cystine deprivation) (Figure 6C and D). These results suggest that PaLung cells are more resistant to ferroptosis compared to WI-38 cells.

**Figure 6.**
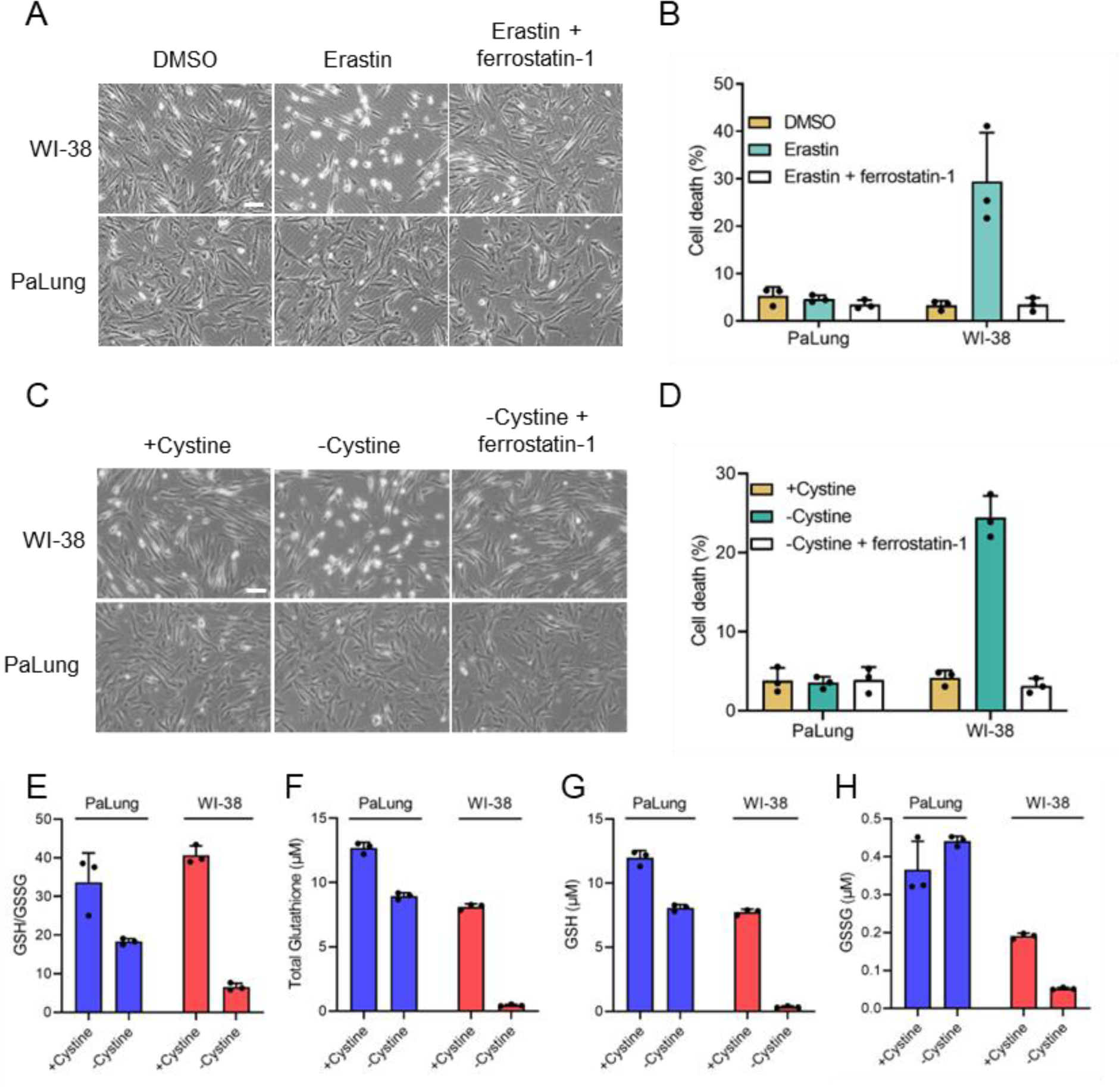
PaLung cells display high resistance to ferroptosis. (A-B) WI-38 or PaLung cells were treated with 2.5 µM erastin and/or 1 µM ferrostatin-1. Representative images were taken at 6 hours using phase-contrast microscopy (A). Propidium iodide (PI) exclusion assay was performed 24 hours after erastin treatment (B). (C-D) WI-38 or PaLung cells were cultured in media with or without cystine. Ferrostatin-1 (1 µM) was treated simultaneously. Representative images were taken at 8 hours using phase-contrast microscopy (C). PI exclusion assay was performed at 24 hours after cystine deprivation (D). (E-H) WI-38 or PaLung cells were cultured in media with or without cystine for 6 hours. Intracellular glutathione levels were measured. Reduced glutathione (GSH)/oxidized glutathione (GSSG) ratio (E), total glutathione (the sum of GSH and GSSG) (F), GSH (G), and GSSG (H) levels were measured. Scale bars, 50 μm. The mean ± SD of three independent experiments is shown.

To better understand the link between glutathione concentrations and ferroptosis resistance in PaLung cells, we measured the amounts of glutathione in WI-38 and PaLung cells under cystine deprivation. Both WI-38 and PaLung cells showed decreased GSH/GSSG ratio under cystine deprivation, which indicates both cell lines are under oxidative stress (Figure 6E). Cystine deprivation decreased total glutathione levels in WI-38 cells by 94.5%, nearly depleting glutathione reserves. In contrast, cystine deprivation in PaLung cells resulted in only a modest 29.5% decrease, and PaLung cells still maintained a total glutathione level higher than non-cystine-deprived WI-38 cells. The higher levels of total glutathione, GSH, and GSSG under cystine deprivation might explain the high resistance of Palung cells under ferroptosis-inducing conditions (Figure 6F, G, and H).

## Methods and materials

### Cell cultures and reagents

WI-38 cells (*H. sapiens*) were purchased from Coriell Institute. PaLung cells (*P. alecto* lung-derived cell line) were established as previously described (Koh *et al*, 2019; Crameri *et al*, 2009). All the cell lines were cultured in high-glucose Dulbecco’s modified Eagle’s medium (DMEM) (#11965, Gibco, Life Technologies) supplemented with 10% FBS (HyClone, GE Healthcare Life Science), penicillin (100 units/ml) and streptomycin (100 mM/ml; Gibco, Life Technologies) in 5% CO_2_-humidified atmosphere at 37°C. For glucose deprivation, cells were washed with phosphate-buffered saline (PBS) three times and cultured in glucose-free DMEM (#10966, Gibco, Life Technologies) with 10% dialyzed FBS. For cystine deprivation, cells were washed with PBS and cultured in cystine, methionine, glutamine-free DMEM (#21013024, Gibco, Life Technologies) supplemented with 0.2 mM L-methionine, 4 mM L-glutamine, and 10% dialyzed FBS. L-glutamine was purchased from Invitrogen. L-methionine and L-cystine were kindly provided by Dr. Jean-Paul Kovalik (Duke-NUS Medical School, Singapore). Oligomycin, FCCP, rotenone, antimycin A, 2-deoxyglucose, and erastin were purchased from Sigma-Aldrich. Ferrostatin-1 was purchased from Med Chem Express.

### Propidium iodide (PI) exclusion assay

Cells were stained with PI to determine the percentage of cell death. Media containing floating cells were collected, combined with trypsinized cells, and centrifuged. The cell pellet was washed once with PBS. After centrifugation, cells were resuspended and stained with PI (10 μg/mL) for 10 mins at room temperature. Data were collected with MACSQuant analyzer (Miltenyi Biotec). Quantification and analysis of the data were done with Flowjo software.

### Transcriptomics (sample preparation and initial bioinformatics)

RNA was extracted from WI-38 and PaLung cells using RNeasy Plus Mini Kit (Qiagen). 1000 ng of total RNA from each sample was used to generate RNA-seq libraries using TruSeq Stranded Total RNA Library Prep Gold according to the manufacturer’s instructions (Illumina). Library fragment size was determined using DNA1000 Assay on the Agilent Bioanalyzer (Agilent Technologies). 2X150PE sequencing was subsequently performed on the libraries using HiSeq3000 equipment (Illumina). The resulting reads were cleaned/trimmed and demultiplexed, followed by mapping to either the PA genome (annotation 102, 2018) or the hg19 genome using cufflinks/Tophat.

### Transcriptomics, DEseq, GeTMM, and GSEA

For differential expression (DE) analysis, the raw read counts were input to the R package DESeq2 (Love *et al*, 2014). Genes with counts per million (CPM) < 1 in more than 3 out of 6 samples (3 from PaLung and 3 from WI-38) were discarded from downstream analysis. To account for differences in transcript length between the two species, the individual transcript lengths were supplied as an additional normalization factor to DESeq2. Since the inter-species analysis of RNAseq data does not have conventional workflows, we repeated DE gene identification using an alternative workflow with Gene length corrected Trimmed Mean of M-values (GeTMM) (Smid *et al*, 2018). This method uses RPKM as input and hence accounts automatically for differences in transcript lengths. This workflow was implemented using the R package EdgeR (Robinson *et al*, 2010). For Gene set enrichment analysis (GSEA), we used the GSEApreranked module in GSEA version 4.1.0 (Subramanian *et al*, 2005), using the DESeq2 results as input. For ranking genes, we used the π-value metric [*LFC\** (−*log*_10_(*pvalue*))] (LFC = Log_2_ fold change) (Xiao *et al*, 2014). GSEA was run against the complete Gene ontology biological process (GO BP) gene set list (containing 18356 gene sets).

### Mitochondria isolation

Mitochondria were isolated from WI-38 and PaLung cells using mitochondrial isolation kit from Miltenyi Biotech as described in the manufacturers’ protocols. In brief, 1×10^7^ cells from WI-38 and PaLung cells were lysed in the ice-cold hypotonic lysis buffer for 60 seconds followed by mechanical disruption of cell membrane using a mini homogeniser pestle gun for 60 seconds. The suspension was centrifuged at 700 x g for 5 minutes at 4°C and supernatant was collected. The supernatant containing the mitochondria was incubated with the TOM-22 antibody conjugated with MACS magnetic beads (Miltenyi Biotech) and pulled down using the magnetic columns. The mitochondria fractions were eluted with 0.1 M glycine pH3.5, neutralized with Tris-HCl pH7.5, and stored at 80°C until the time of proteomics profiling.

### Proteomics profiling and analysis

The mitochondrial fractions were lysed in 8 M urea pH 8.5 and incubated with 20 mM Tris(2-carboxyethyl)phosphine hydrochloride (TCEP) for 20 min at 25°C followed by alkylation with 55 mM chloroacetamide at 25°C for 30 min. Proteins were digested with trypsin overnight at 25°C. Digested peptides were acidified with 1% trifluoroacetic acid, desalted on C18 plates Oasis (Waters), and labelled with TMT sixplex reagent (Thermo Scientific) according to the manufacturer protocol. Labelled samples were further fractionated with high pH reverse phase using spin columns packed in-house, and five fractions were collected: 10%, 17.5%, 25%, 30%, and 50%. The fractions were separated on a 50 cm x 75 µm Easy-Spray column using Easy-nLC system coupled with an Orbitrap Fusion Tribrid mass spectrometer (Thermo Scientific). The LC-MS/MS parameters for Fusion: peptides were separated over a 120 min gradient, using mobile phase A (0.1% formic acid in water) and mobile phase B (0.1% formic acid in 99% acetonitrile), and eluted at a constant flow rate of 300 nl/min. Acquisition parameters were as follows: data dependent acquisition (DDA) with survey scan of 60,000 resolution, AGC target of 4 × 10^5^, and maximum injection time (OT) of 100 ms; MS/MS collision induced dissociation in Orbitrap 15,000 resolution, AGC target of 1 × 10^5^, and maximum IT of 120 ms; collision energy NCE=35, isolation window 1.0 m/z. Peak lists were generated in Proteome Discoverer 2.1, and a search was done using Sequest HT (Thermo Scientific) with human Uniprot and fruit bat Uniprot databases. The following search parameters were used: 10 ppm MS; 0.06 Da for MS/MS with following modifications: Oxidation (M) Deamidation (N,Q), TMT adduct (N-term, K) Carbamidomethyl (C). Peptides detected in WI-38 samples were automatically mapped to their source human protein Uniprot ID. To map the peptides detected in PaLung samples to the corresponding *P. alecto* Uniprot ID, we used a two-step approach. First, we used inParanoid (O’Brien *et al*, 2005) to identify known orthologs between *H. sapiens* and *P. alecto* species. In cases where orthologs were not available, we used blast using the peptide sequence as a query to detect the possible source of *P. alecto* protein. Protein abundances were obtained by summing the abundances of peptides derived from them. We identified differentially expressed proteins by first performing median normalization on all samples from WI-38 and PaLung cells (total 6 samples), followed by a student’s t-test for all proteins abundances. P-values were corrected using the false discovery rate method of Benjamini-Hochberg.

For the mitochondrial protein-protein interaction network in Figure 2C, we first obtained a list of all known mitochondrial proteins from the Mitocarta (Calvo *et al*, 2016) and IMPI databases. These were then input into STRING (Szklarczyk *et al*, 2019) to identify all high-confidence pairwise protein-protein interactions. We used Cytoscape (Shannon *et al*, 2003) to both visualize the network and overlay fold change values of detected proteins onto the network. Clustering of the network and Reactome enrichment of identified sub-networks were performed using the ClusterOne app of Cytoscape. For Gene set enrichment analysis (GSEA), the raw abundances of all samples were used as input and GSEA was run against the GO BB gene set list, as with RNAseq data.

### Metabolomics

WI-38 and PaLung cells were harvested according to the protocol outlined in the document ACB.1.0.0 provided by Human Metabolome Technologies (HMT Japan). Targeted quantitative analysis was performed by HMT, using capillary electrophoresis mass spectrometry (CE-TOFMS and CE-QqQMS). Absolute abundances (adjusted for cell numbers) were obtained for a total of 116 metabolites (54 and 62 metabolites in the cation and anion modes, respectively)

### Computational Flux analysis

#### Metabolic network reconstruction

A network model of central metabolism in humans was obtained from Mitocore (Smith *et al*, 2017) (485 reactions, 371 metabolites). This network contains most mitochondrial reactions and pathways, with additional reactions for glycolysis, pentose phosphate pathway, and import and export of amino acids, ions, and other metabolites. The mitocore model also contains a list of genes that catalyze each reaction in a gene-reaction rules table (Both mandatory and optional genes). Using this base model, separate context-specific reconstructions for PaLung and WI-38 cell lines were obtained manually using the proteomic and transcriptomic data as follows. The expression levels of all enzymes of the mitocore model were checked in our proteomics and RNAseq datasets, and enzymes were marked as missing if they had < 5 counts in 2 out of 3 samples (for each PaLung and WI-38) for transcriptomics, and < 2000 abundance in 2 out of 3 samples for proteomics. Reactions in the mitocore model whose activity depended on the presence of missing genes were iteratively removed from the model while ensuring that the removal of reactions would not result in absence of flux-carrying capacity for ATP production, TCA cycle, or glycolysis. The final reconstructed model for PaLung cells contained 409 reactions and 324 metabolites, and the reconstructed model for WI-38 cells contained 437 reactions and 341 metabolites. We call these PaLung and WI-38 models the “reconstructed models”.

#### Flux sampling

For each species, we first obtained a feasible range of metabolic flux levels for each reaction by performing flux sampling on the “reconstructed” model for that species. To perform flux sampling, we generated 5000 flux vectors through uniform sampling using the COBRA toolbox v3.0 (Heirendt *et al*, 2019). We call this round of flux sampling the “control simulation” as there are no constraints imposed on the flux of the reactions for Complex I and oxygen consumption, meaning that these reactions are free to assume any flux values that satisfy the steady-state assumption. Next, we repeated the flux sampling process for a case where the PaLung model was forced to have higher flux through the Complex I reaction than the WI-38 model, and lower flux through the oxygen consumption reaction than the WI-38 model. We call this the “constrained simulation.” The setting of constraints is explained in the following section.

#### Constraints for flux sampling

Constraining reaction fluxes in a metabolic network is usually accomplished by setting a lower bound or an upper bound (or both) on the feasible range of the flux values for individual reactions. When a lower bound is set, flux sampling will only output flux vectors where the flux for that reaction is greater than or equal to the set lower bound. Similarly, setting an upper bound ensures that flux sampling will only output flux vectors where the reaction flux is less than or equal to the set upper bound. To ensure that the PaLung model would have higher Complex I activity and lower mitochondrial oxygen consumption than the WI-38 model, we set flux constraints as follows. We first computed the maximum possible fluxes of the Complex I reaction (Reaction ID: CI_MitoCore) in both PaLung and WI-38 models. We then set the lower bound of the PaLung Complex I reaction flux to a value equal to 70% of its theoretical maximum. Similarly, we set the upper bound of the WI-38 Complex I reaction at a value equal to 30% of its theoretical maximum value. This ensured that the PaLung model would have higher flux through the Complex I reaction, in comparison to the WI-38 model. A similar process of identifying maximum possible fluxes and setting lower and upper bounds was followed for the oxygen consumption reaction (Reaction ID: O2tm) to ensure that the lower bound of the reaction in the WI-38 model was higher than the upper bound in the PaLung model. Flux sampling was then performed for both the constrained PaLung model and the constrained WI-38 model to obtain 5000 flux vectors (as with the control simulation).

### Mitosox assay and antimycin A treatment

PaLung and WI-38 cells were cultured on 6-well plates and treated with either DMSO or antimycin A (1 μΜ) for 16 hours. Antimycin A treated cells were washed twice with PBS before adding fresh media containing 2.5 μΜ MitoSox Red. Cells were stained with MitoSox Red for 60 minutes and subsequently collected for flow cytometry. Quantification and analysis of the data were performed by Flowjo software.

### NADPH/NADP+ measurement assay

Intracellular NADPH/NADP+ were measured using the NADP/NADPH Quantification Kit (ab65349, Abcam) according to the manufacturer’s instructions. Briefly, 6×10^5^ cells were lysed with 350 μL of extraction buffer. For the reaction, 50 μL of the final sample was used. Signal intensities for NADPH were examined by OD measurements at 450 nm using Infinite M200 plate reader (TECAN).

### GSH/GSSG measurement assay

Intracellular GSH/GSSG was measured using GSH/GSSG-GloTM luminescent assay (Promega) according to the manufacturer’s instructions. Briefly, 2 x10^4^ cells in 96-well plates were lysed in the indicated condition.

### Mitochondrial oxygen consumption rate (OCR) measurement

OCR was measured using a Seahorse Bioscience XF96 Extracellular Flux Analyzer (Seahorse Bioscience). 2 x 10^4^ cells were plated into Seahorse tissue culture 96-well plates. Cells were cultured in Seahorse assay media containing 10 mM glucose and 2 mM glutamine and incubated in a CO2-free incubator for an hour before measurement. XF Cell Mito Stress Test Kit was used to analyze mitochondrial metabolic parameters by measuring OCR. Oligomycin (1 μΜ) was injected to determine the oligomycin-independent lack of the OCR. The mitochondrial uncoupler FCCP (1 μΜ) was injected to determine the maximum respiratory capacity. Rotenone (1 μΜ) and antimycin A (1 μΜ) were injected to block Complex I and Complex III of the electron transport chain.

### Extracellular acidification rate (ECAR) measurement

ECAR was measured using a Seahorse Bioscience XF96 Extracellular Flux Analyzer (Seahorse Bioscience). 2 x 10^4^ cells were plated into Seahorse tissue culture 96-well plates. Cells were cultured in Seahorse assay media containing 10 mM glucose and 2 mM glutamine and incubated in a CO2-free incubator for an hour before measurement. XF Cell Glycolysis Stress Test Kit was used to analyze glycolytic metabolic parameters by measuring ECAR. Glucose (10 mM), oligomycin (1 μΜ), and 2-deoxy-D-glucose (2-DG; 50 mM) were injected sequentially.

## Discussion

In this study, we integrated high throughput-omics and computational metabolic modeling to perform a novel comparison of central metabolism between cell lines from two mammalian species. Specifically, we have identified core differences in mitochondrial metabolism between primary lung fibroblasts of the black flying fox fruit bat *P. alecto* (PaLung) and the primary human lung fibroblast cell line WI-38. Although data are still limited to these two cell lines, this is the first comprehensive analysis combining proteomics, transcriptomics, metabolomics, and constraint-based flux modelling between human and bat cells. Our analysis suggests that PaLung cells exhibit basal metabolism that resembles an ischemic-like state and that this state may be linked to low or reverse activity of Complex II of the electron transport chain (also called succinate dehydrogenase, SDH). Compared to WI-38 cells, PaLung cells also show a higher tolerance to cellular stresses such as nutrient deprivation, and ischemia/ROS-driven cell death via ferroptosis.

Hypoxia response (including glycolysis) and OxPhos compensate for each other by gene expression, dependent on different levels of oxygen availability. However, our analyses (GSEA from whole-cell transcriptomics) indicated that genes related to glycolysis and OxPhos were simultaneously upregulated in PaLung cells (bat) compared to WI-38 cells (human). Although multiple glycolytic genes were upregulated in PaLung cells (Supplementary Figure S2), lactate production (inferred via both LDH expression levels and ECAR measurement) in PaLung cells was not higher than in WI-38 cells. This raises an important caveat about using extracellular acidification rate (ECAR) to infer glycolytic flux, because ECAR depends only on the amount of lactate produced, and glycolytic pyruvate that enters the TCA cycle may remain invisible to ECAR measurements.

Our observation about the upregulation of OxPhos in PaLung cells via GSEA (transcriptomics and proteomics) also raises an important point of concern, namely that high signals of a certain pathway in gene set enrichment analysis (GSEA) can be driven by a narrower subset of genes. Hence, while interpreting GSEA results, it is necessary to survey the constituent genes that are responsible for the predicted up/down regulation of the overall pathway. Indeed, we found that the upregulation of OxPhos pathway in PaLung cells in GSEA was due to high gene expression of the Complex I components in the ETC. However, our experimental analysis revealed a puzzling result: despite high Complex I expression, oxygen consumption was low in PaLung cells.

To interpret the metabolic implications of heightened Complex I and lower oxygen consumption, we turned to constrained-based metabolic flux sampling. Flux sampling is a technique that can simulate possible states of a metabolic network. Compared to higher resolution methods such as isotope-labeled fluxomics, flux sampling is a coarse-grained qualitative approach to study metabolic flux. However, it can be used as a tool to generate testable hypotheses in exploratory studies such as ours. Indeed, metabolomics was able to verify our simulation predictions of succinate accumulation in PaLung cells, resulting from low/reverse activity (fumarate to succinate) of Complex II of ETC (succinate dehydrogenase). The succinate-to-fumarate ratio in well-oxygenated PaLung cells was found to be similar to those found in ischemic states of human cells from multiple tissues (Chouchani *et al*, 2014). From this, we infer that the TCA cycle in PaLung cells does something different than the TCA cycle in human WI-38 cells, which is also corroborated by our observation that PaLung cells have a low metabolic rate. We also confirmed using cell culture assays that PaLung cells have a much longer survival during glucose deprivation than WI-38 cells.

We also found that PaLung cells had higher loads of mitochondrial ROS, consistent with having the lower expression of *SOD1* and *SOD2.* On the other hand, PaLung cells highly express glutathione peroxidase 3 (*GPX3*), contain 2-fold higher levels of glutathione, and maintain a higher NADPH/NADP ratio compared to WI-38 cells. These data suggest that PaLung cells maintain a standing pool of NADPH and glutathione that can act against high levels of mitochondrial ROS. Therefore, the previously published generalization that bats have lower free radical production than other mammals (Brunet-Rossinni, 2004; Brown *et al*, 2009) may require moderation. Future work can address whether PaLung cells have a higher threshold for homeostatic ROS and whether ROS have both beneficial and detrimental effects in bats, as has been observed in other organisms (Clément & Pervaiz, 2001; Shields *et al*, 2021).

The key novelty of our results is the independent identification of a basal ischemic-like state in PaLung cells accompanied by higher ROS production and higher glutathione and NADPH levels. Multiple studies have shown strong links between ischemia, ROS, and ferroptosis (an iron-dependent, non-apoptotic form of cell death). Despite having ischemic-like metabolism and higher ROS levels, we observed that PaLung cells were more resistant to ferroptosis than WI-38 cells. Ferroptosis occurs via ROS-induced lipid peroxidation following ischemia and is conventionally blocked via the action of glutathione peroxidase 4 (*GPX4*). While *GPX4* expression levels were low in PaLung cells, *GPX3* (a commonly secreted form of glutathione peroxidase) was highly expressed. Future studies are needed to determine the function of GPX3 in the stress response and whether it contributes to the ferroptosis-resistance of PaLung cells.

Our identification of a metabolic state in bats that resembles ischemia is also tied to conventional wisdom about bat flight. During flight, the metabolic rate of bats can go 2.5-3 times higher than that of other mammals, consuming approx. 1200 calories per hour, resulting in an immense drain on stored energy reserves, and the potential depletion of up to 50% of stored energy in fructivore bats (Thomas, 1975; Voigt & Speakman, 2007; Kelm *et al*, 2011). Thus, a basal state with a low metabolic rate may serve to conserve energy during non-flying periods. In addition, the accumulated succinate during this ischemic-like state can also be transported into the bloodstream and serve as a metabolic stimulant for other fast-metabolizing cell types as needed, similar to examples of succinate shuttling between two cell types in retinal tissues (Bisbach *et al*, 2020). Low metabolic rates have also been linked to longevity by many studies, which demonstrate that calorie restriction contributes to an extended lifespan across various species. In our experiments, *P. alecto* also showed higher expression of genes involved in NAD synthesis and its phosphorylation pathways, in agreement with other studies that have linked higher flux through the NAD biosynthesis pathway with slower ageing (Chini *et al*, 2017).

A limitation of our study is that we performed multi-omics comparisons between individual cell lines. Extending the current study to primary tissues or cell lines from other species of bats might provide further information about generalized metabolic differences between bats and humans. The current study can be considered a starting point to both generate hypotheses for future work and to establish analysis pipelines for inter-species comparisons of metabolism, using multi-omics data and computational flux modeling simulations.

In summary, using multi-omics datasets (transcriptomic, proteomic, metabolomic) and computational modeling, we have compared the basal metabolic states of a human cell line and a *P. alecto* cell line. Our work points toward important differences in ETC regulation (low/reverse Complex II activity) and antioxidant response (higher glutathione and NADPH) that could contribute to ischemia/redox management and ferroptosis resistance in *P. alecto*. Future research can extend the idea further to identify if such regulation is a pan-bat feature and if such metabolic fingerprints might also contribute to cancer resistance and increased longevity in bats.

## Acknowledgments

The authors thank Dr. Akshamal M. Gamage and Prof. Lena Ho for their insightful commentary on the manuscript.

## Data availability

Transcriptomic data are deposited in the NCBI GEO database (ID: GSE215934). Matlab script used for flux sampling can be found at https://github.com/nsuhasj/PalungWI38FluxSim. Other data supporting the findings of this study are available within the article and its supplementary materials.

## Author Contributions

NSJ, JK, YI, KI, LFW, and LTK conceptualized the project. JK, YI, and ATI performed the transcriptomics experiments; JK, YI, and RMS performed the proteomics experiments; YL and YI performed the OCR, ECAR, glucose deprivation, and ferroptosis experiments; JK and YI performed all other assays; NSJ, ATI, and LTK performed the statistical analysis and modeling; NSJ and YL performed all visualization; NSJ, JK, YL, YI, KI, and LTK wrote the manuscript.

---

**Figure S1.**
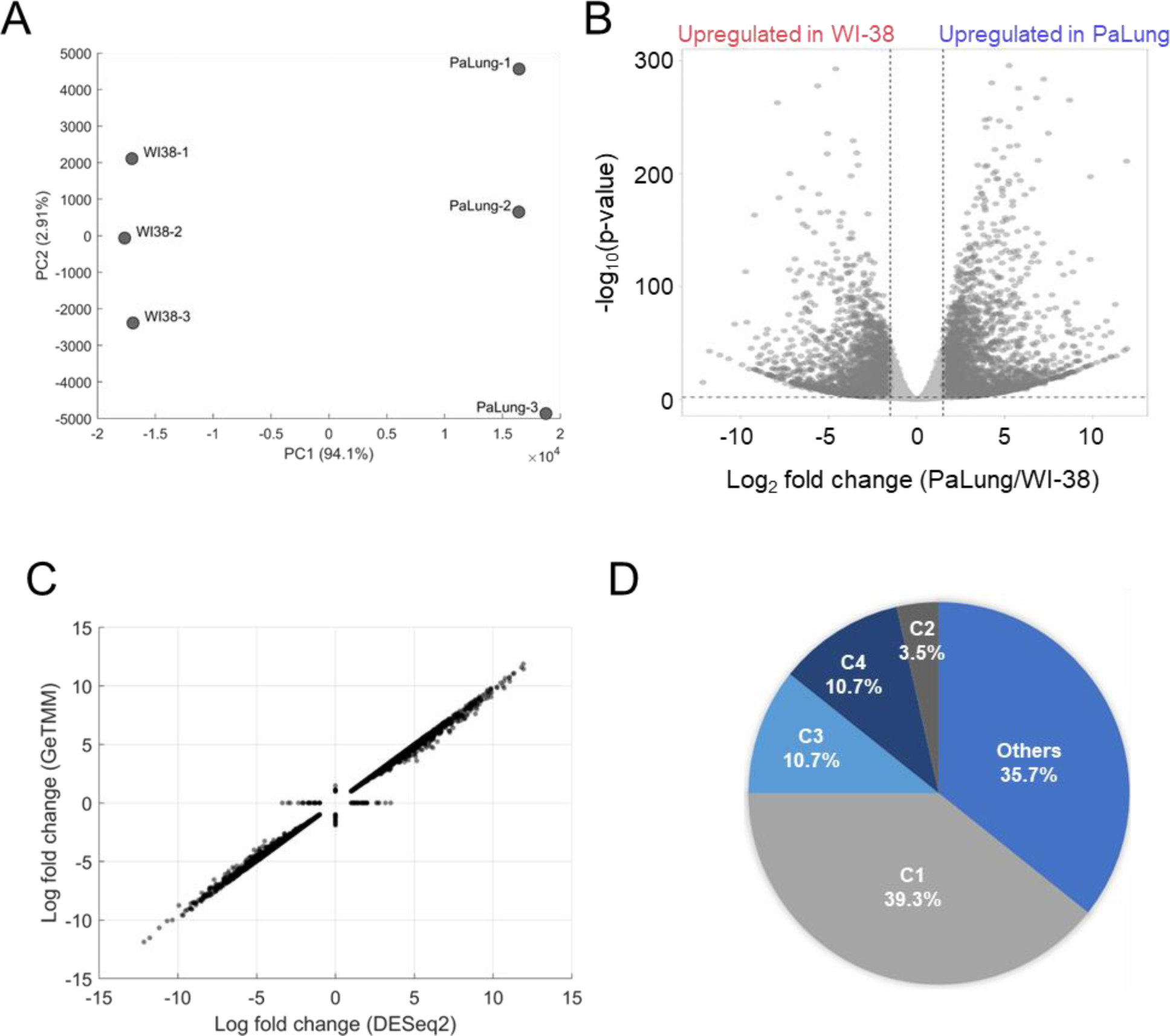
Analysis of Transcriptomics data from PaLung and WI-38 cells. (A) Principal Component Analysis plots showing the separation of the PaLung and WI-38 transcriptomics datasets. (B) Volcano plot showing the differentially expressed (DE) genes (|Log2 Fold Change| >= 1 and FDR < 0.05). (C) Correlation between the DE gene fold changes estimated via the DESeq2 pipeline and the GeTMM pipeline (see Methods). The points on the X-axis represent LFC of genes that were considered differentially expressed in DESeq2 but not in GeTMM. Conversely, points on the Y-axis represent genes considered DE in the GeTMM pipeline but not in the DESeq2 pipeline. (D) Pie chart showing that the GSEA-suggested upregulation of ETC in transcriptomics data is a result dominated by genes corresponding to Complex I subunits. C1 = Complex I; C2 = Complex II; C3 = Complex III; C4 = Complex IV of the ETC.

**Figure S2.**
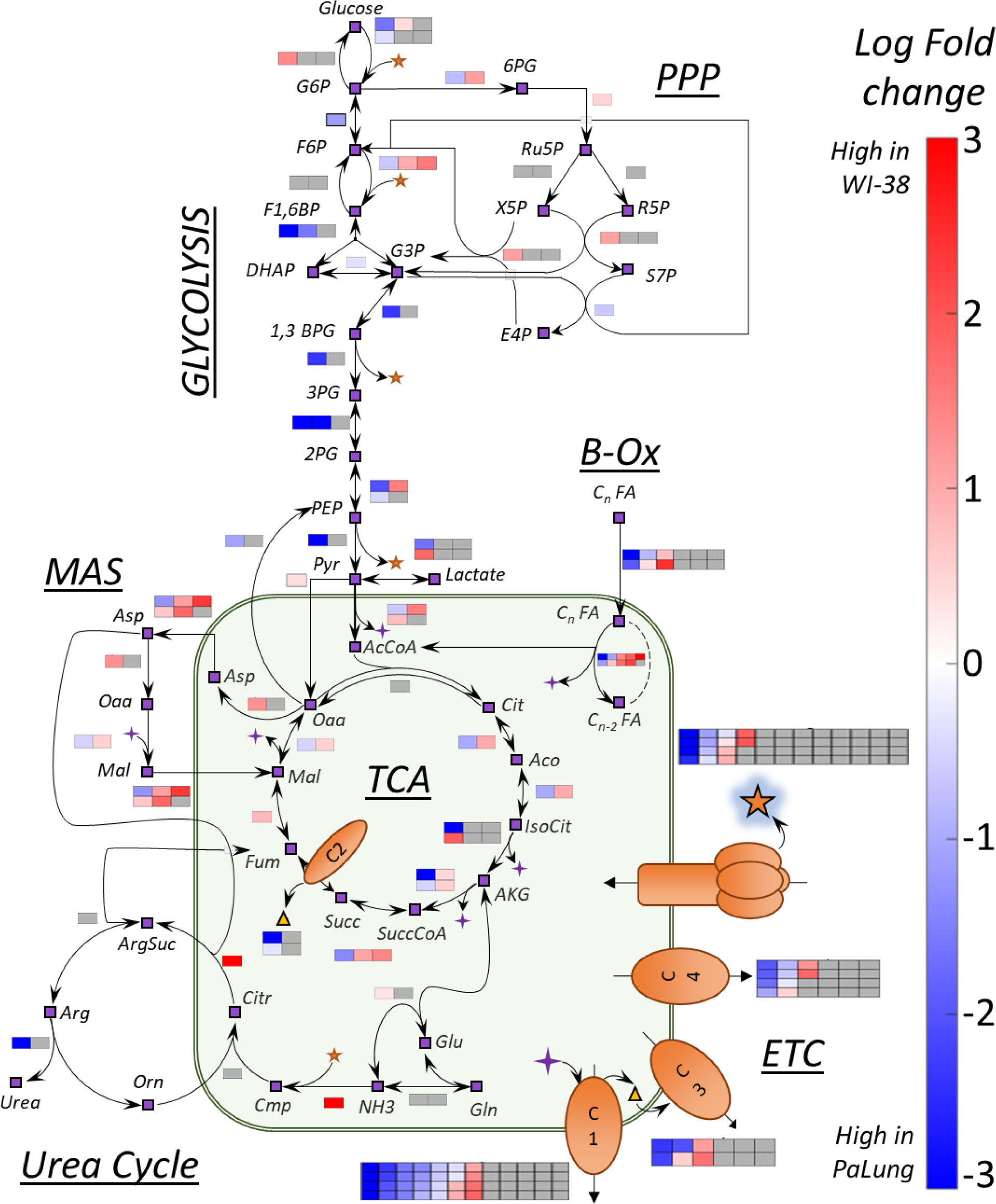
A summary of transcriptomics Log fold changes overlaid onto key metabolic reactions from central carbon metabolism. Shown here is a network of reactions in central carbon metabolism that includes reactions from multiple pathways (glycolysis, pentose phosphate pathway, TCA cycle, fatty acid oxidation, electron transport chain, urea cycle and malate aspartate shuttle). The nodes of the network indicate metabolites and are shown as purple boxes with black outlines. The black arrows represent metabolic reactions. The colored boxes adjacent to the arrows represent the log fold change (LFC) of a gene involved in catalyzing the specific metabolic reaction (subunit/isoform/alternate genes). The color scale indicates the extent to which the gene is upregulated in PaLung (blue) or WI-38 (red). Grey squares indicate that a particular gene/transcript implicated in the reaction was either not detected or was not differentially expressed between PaLung and WI-38 samples.

**Figure S3.**
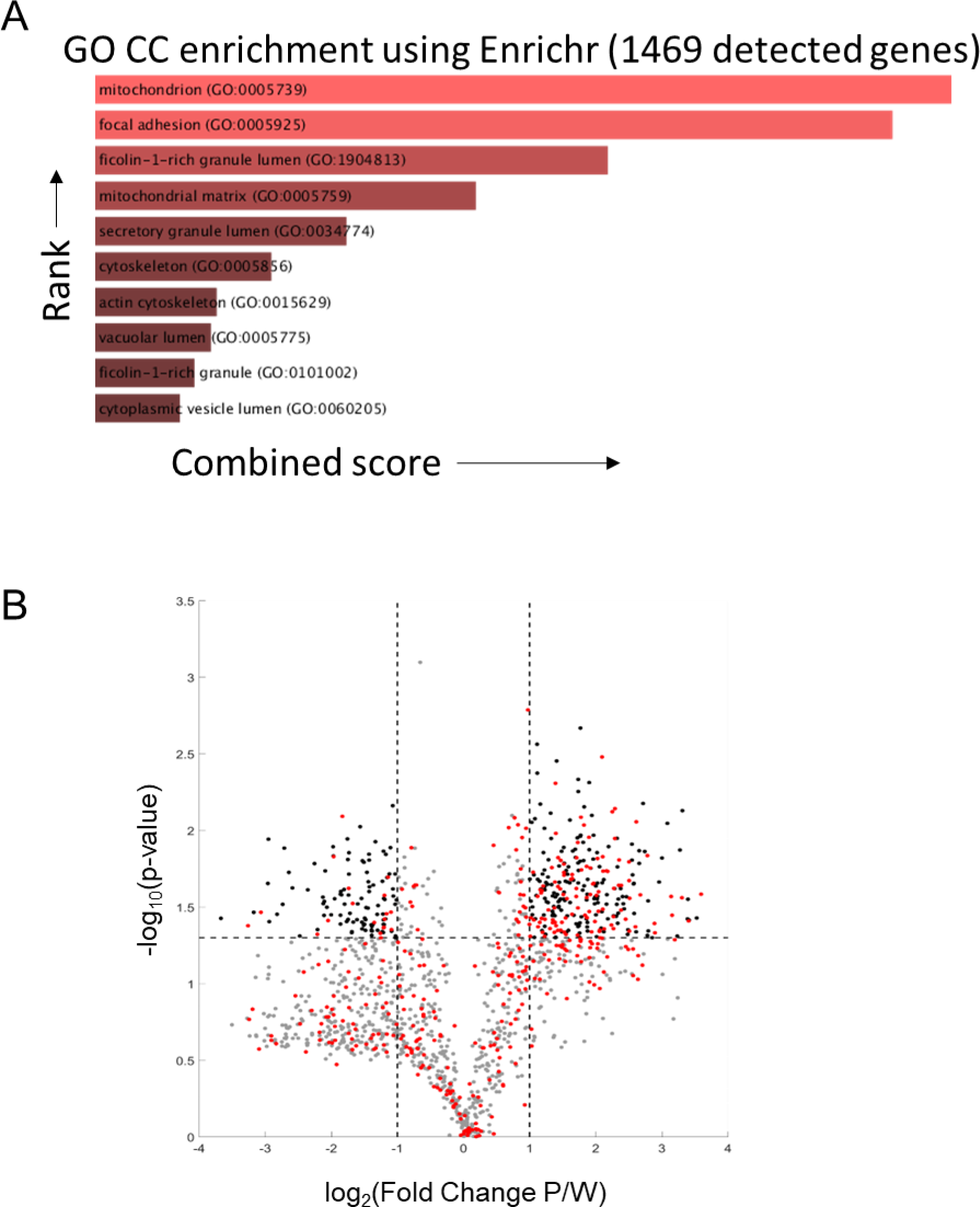
Proteomic analysis of PaLung and WI-38 data. (A) Gene ontology category Cellular (GO CC) enrichment using the Enrichr tool for compartmental enrichment, on the 1469 genes detected in the proteomics dataset shows enrichment for the mitochondrial compartment. (B) Volcano plot showing the differentially expressed (DE) proteins (|Log2 Fold Change| >= 1 and FDR < 0.05). Grey dots represent non-mitochondrial proteins that are not differentially expressed. Black dots show non-mitochondrial proteins that are differentially expressed. Red dots show all mitochondrial proteins.

**Figure S4.**
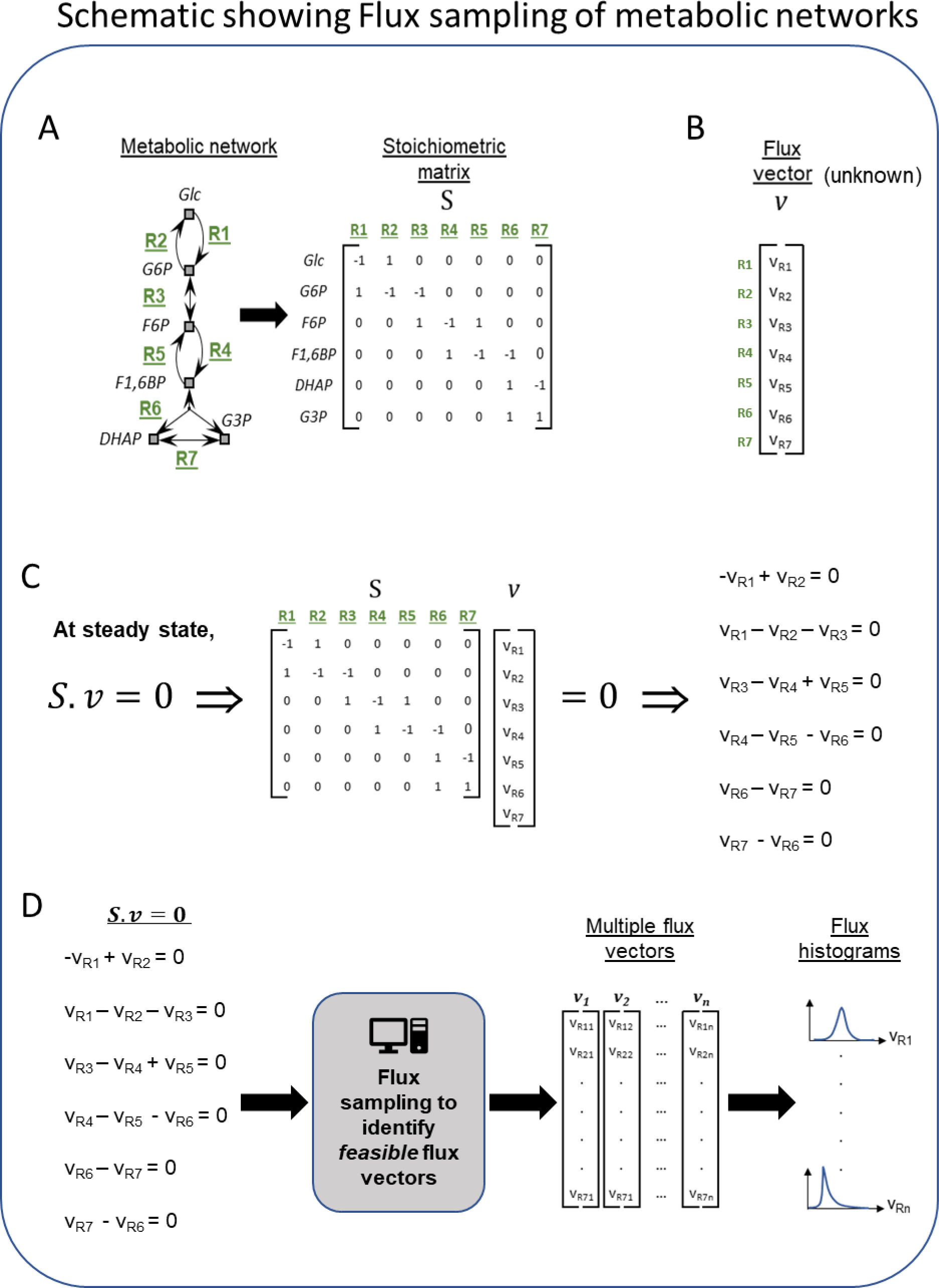
Schematic of the workflow for metabolic flux modeling. (A) Left is a toy network of metabolic reactions showing a few reactions from the glycolysis pathway for illustration. This network of metabolic reactions (defined by nodes and arrows) is converted from the format of reaction equations into the format of a matrix called the stoichiometric matrix. Each row of the matrix represents a single metabolite in the network and each column represents a single reaction in the network. The (*i,j*) element of matrix *S* represents the stoichiometric coefficient of metabolite *i* in reaction *j*. Stoichiometric coefficients indicate how many copies of each reactant or product are consumed or produced when the reaction is utilized (often 1 or −1). By converting all the reactions of a metabolic network into this matrix format, a single column represents a single reaction in the metabolic network, including all metabolites produced or consumed in it. For example, the first column of the matrix (corresponding to reaction R1) has the value of −1 for the *Glc* row and 1 for *G6P* row, indicating that in reaction R1, one unit of *Glc* is consumed and one unit of *G6P* is produced. Likewise, each row of the matrix represents all sources of production and consumption for a single metabolite. For example, the second row of the matrix (corresponding to the metabolite *G6P*) has the values of 1, −1, and −1 for the reactions R1, R2, and R3 respectively. This indicates that one unit of *G6P* is produced in reaction R1, while one unit of G6P is consumed in reactions R2 and R3, respectively. (B) Example of a flux vector. A flux vector contains as many elements as the number of reactions in the network, and their values are typically unknown. Flux values indicate the rate at which the corresponding reaction is utilized (a forward reaction is a positive flux, and a reverse reaction is a negative flux). The goal of flux sampling is to obtain values (or distributions) for each element of the flux vector. (C) The product of the stoichiometric matrix and the flux vector (written as the product *Sv)* results in a set of linear algebra equations. Each linear equation has flux variables and stoichiometric coefficients. If we assume the flux is balanced in the system, then everything produced has somewhere to go or some reaction to consume it. This steady-state assumption is described mathematically by setting the matrix product to zero (*Sv* = 0). (D) The process of flux sampling searches the space of possible flux vectors *v* to find those that satisfy *Sv* = 0. Sampling is typically necessary because the set of equations from (C) is usually underdetermined and does not have a unique solution for the flux variables *v*. The computational sampling process outputs flux vectors that satisfy the equations (i.e., *feasible* vectors), and that also capture a diversity of feasible behaviors for how the metabolic network could achieve steady-state. From this set of flux vectors, we plot histograms (distributions) of the feasible values of the flux variable for each reaction in the metabolic network. This theoretical delineation of feasible behaviors can be used for drawing biological inferences by assuming that the actual biological pathway activity falls within the feasible range of the theoretical model. When comparing two models such as bat and human, inferences arise from reactions or pathways where feasible behaviors become infeasible or vice versa. This can be seen by identifying reactions, whose flux histograms show large differences between the human and bat models.

**Figure S5.**
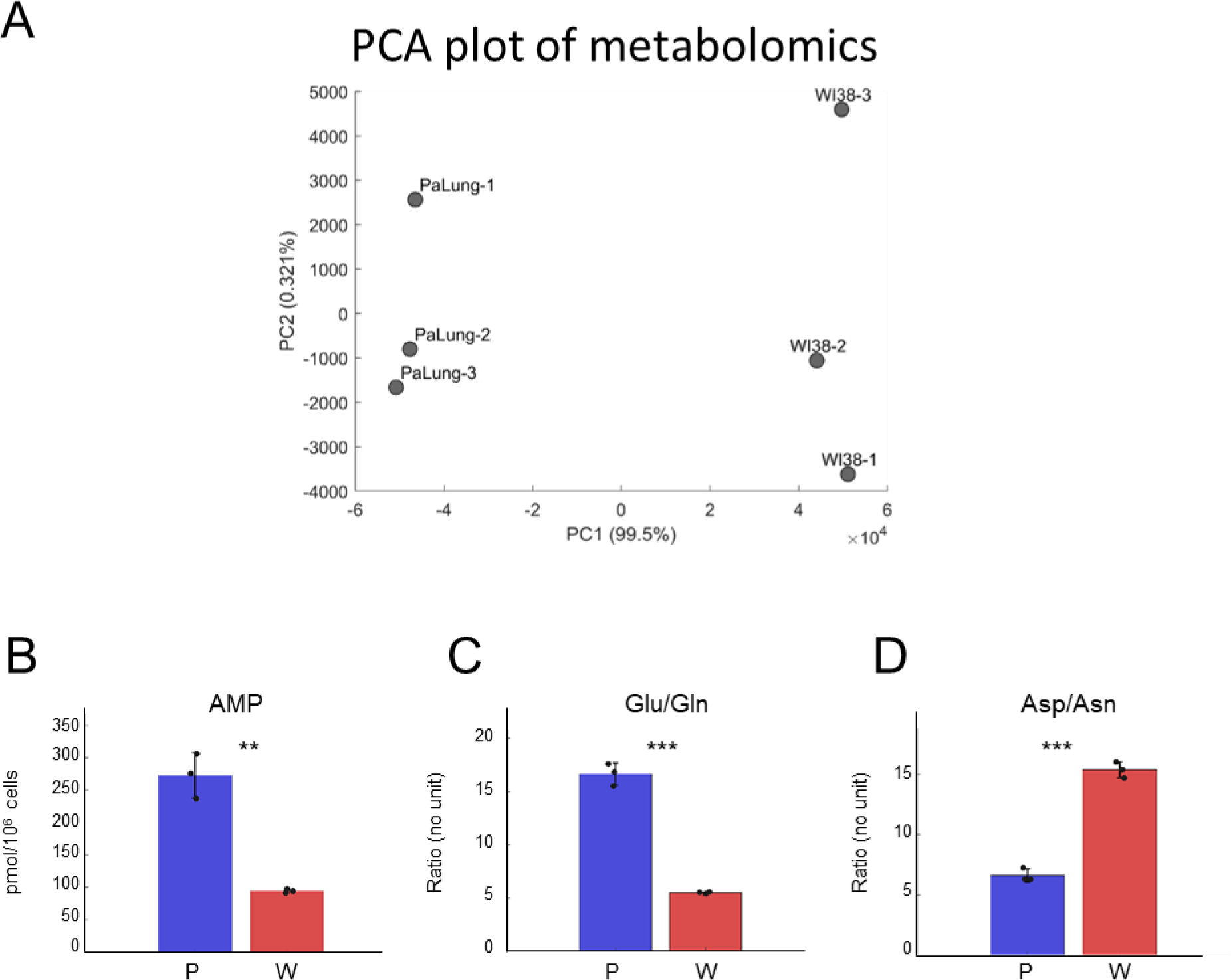
Analysis of the metabolomics data from PaLung and WI-38 cells. (A) Principal Component Analysis (PCA) plot shows the separation between the three PaLung samples and the three WI-38 samples in the metabolomics data. (B-D) Bar plots of AMP (B), the ratios of Glu/Gln (C), and Asp/Asn (D) in PaLung cells (P, blue) and WI-38 cells (W, red) are shown. Bars are the mean ± SD from three independent experiments (n = 3). ** or *** represents p-value ≤0.01 or ≤0.001 respectively (unpaired Student’s two-sided t test). Note that PaLung cells have a higher Glutamate-to-Glutamine ratio and a lower Aspartate-to-asparagine ratio than WI-38 cells. This is also supported by the high expression of the gene ASNS (Asparagine synthetase) in PaLung cells (Supplementary Table 1). ASNS is a cytoplasmic enzyme that can convert aspartate to asparagine concomitantly with the hydrolysis of glutamine to glutamate, allowing for glutamate to be used as an alternative source for energy, or as a precursor metabolite for the synthesis of glutathione.

**Figure S6.**
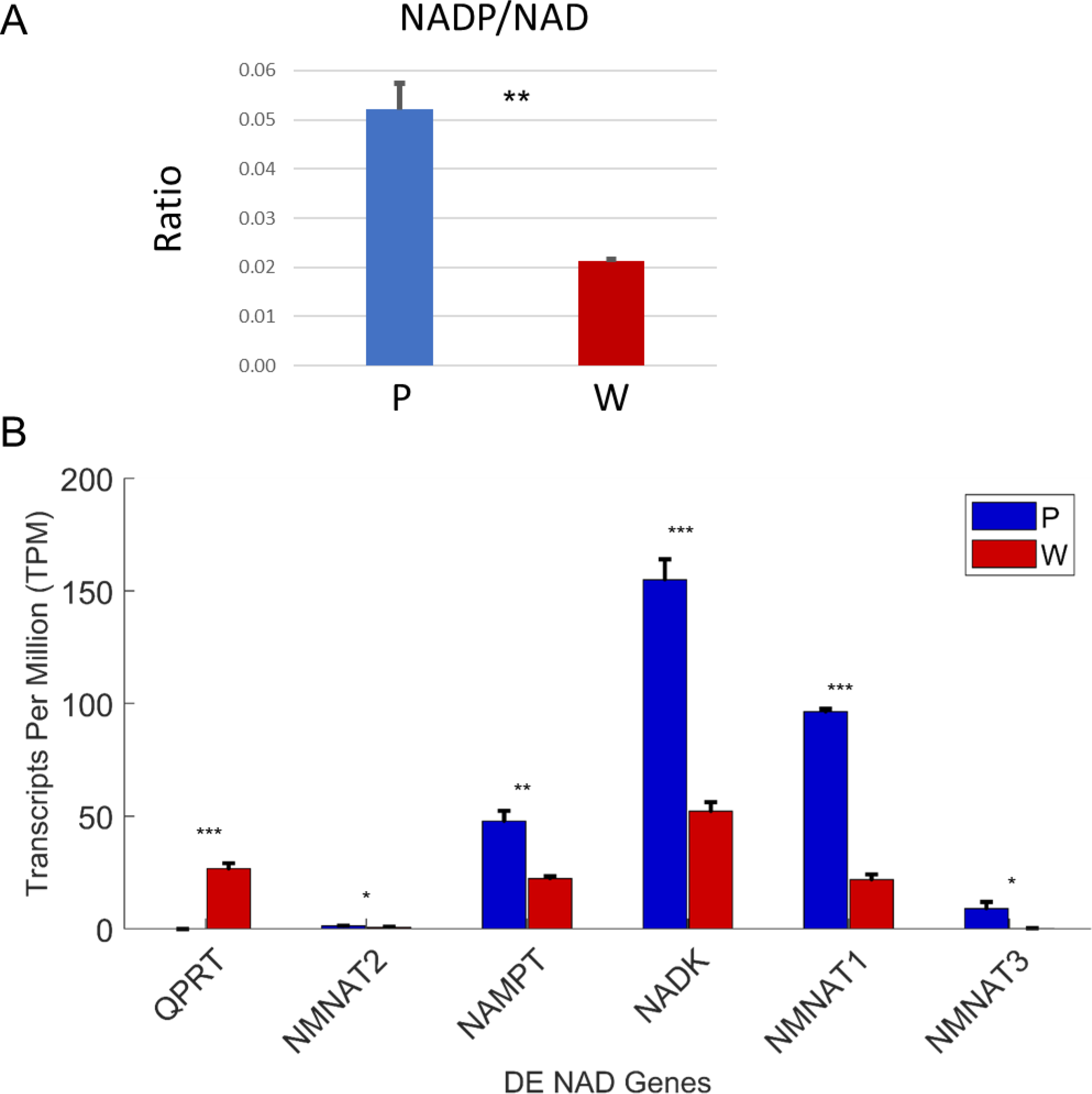
Increased phosphorylation of NAD cofactors in PaLung cells. (A) Ratio of NADP/ NAD in PaLung (P) and WI-38 (W) cells. (B) Bar graphs show the expression levels of the genes involved in NAD synthesis and phosphorylation (as transcripts per million, TPM) in PaLung (blue) and WI-38 (red) cells. *, ** or *** represents p-value ≤0.05, ≤0.01 or ≤0.001 respectively.

